# Consolidated bioprocessing of lignocellulose for production of glucaric acid by an artificial microbial consortium

**DOI:** 10.1101/2020.04.13.039354

**Authors:** Chaofeng Li, Xiaofeng Lin, Xing Ling, Shuo Li, Hao Fang

**Affiliations:** College of Life Sciences, Northwest A&F University, 22 Xinong Road, Yangling 712100, Shaanxi, China; Biomass Energy Center for Arid and Semi-arid Lands, Northwest A&F University, 22 Xinong Road, Yangling 712100, Shaanxi, China

**Author notes:** Corresponding Author. College of Life Sciences, Northwest A&F University, 22 Xinong Road, Yangling 712100, Shaanxi, China. E-mail address (H. Fang).

**Keywords:** Consolidated bioprocessing (CBP), D-glucaric acid, Lignocellulose, Microbial consortium, *Trichoderma reesei*, *Saccharomyces cerevisiae*

## Abstract

The biomanufacturing of D-glucaric acid has been attracted increasing interest and the industrial yeast *Saccharomyces cerevisiae* is regarded as an excellent host for D-glucaric acid production. Here we constructed the biosynthetic pathway of D-glucaric acid in *S. cerevisiae* INVSc1 whose *opi1* was knocked out and obtained two engineered strains, LGA-1 and LGA-C, producing record breaking titers of D-glucaric acid, 9.53 ± 0.46 g/L and 11.21 ± 0.63 g/L D-glucaric acid from 30 g/L glucose and 10.8 g/L *myo*-inositol in the mode of fed-batch fermentation, respectively. Due to the genetic stability and the outperformance in subsequent applications, however, LGA-1 was a preferable strain. As one of the top chemicals from biomass, there have been no reports on D-glucaric acid production from lignocellulose, which is the most abundant renewable on earth. Therefore, the biorefinery processes of lignocellulose for D-glucaric acid production including separated hydrolysis and fermentation (SHF), simultaneous saccharification and fermentation (SSF) and consolidated bioprocessing (CBP) were investigated in this work and CBP by an artificial microbial consortium composed of *Trichoderma reesei* Rut-C30 and *S. cerevisiae* LGA-1 was found to have relatively high D-glucaric acid titers and yields after 7 d fermentation, 0.54 ± 0.12 g/L D-glucaric acid from 15 g/L Avicel, and 0.45 ± 0.06 g/L D-glucaric acid from 15 g/L steam exploded corn stover (SECS), respectively. In attempts to design the microbial consortium for more efficient CBP the team consisted of the two members, *T. reesei* Rut-C30 and *S. cerevisiae* LGA-1, was found to be the best with excellent work distribution and collaboration. This desirable and promising approach for direction production of D-glucaric acid from lignocellulose deserves extensive and in-depth research.

## Introduction

D-Glucaric acid, identified as “top value-added chemical from biomass” by US Department of Energy in 2004 ^1^, is an important platform chemical with a wide variety of applications such as therapeutic uses and biopolymer production ^2, 3, 4^. Conventionally, D-glucaric acid was produced via nitric acid oxidation of D-glucose, a nonselective and expensive process associated with a large exotherm, low yields and toxic byproducts ^4, 5, 6^. Biological production of D-glucaric acid, therefore, has been attracted increasing interest due to the potential for a cheaper and more environmentally friendly process by avoiding costly catalysts and harsh reaction conditions ^3, 4^.

A biosynthetic route from D-glucose to D-glucaric acid consisting of three heterologous genes, myo-inositol-1-phosphate synthase (Ino1) from *Saccharomyces cerevisiae* and *myo*-inositol oxygenase (MIOX) from *Mus musculus*, and urinate dehydrogenase (UDH) from *Pseudomonas syringae*, was constructed in recombinant *Escherichia coli* by Moon et al.^2^ and a low D-glucaric acid titer of 0.72 g/L was reported ^2^. After optimizing the induction and culture conditions, a slight increase in D-glucaric acid titer to 1.13 g/L was achieved ^2, 4^. MIOX was found to be the rate-limiting step of the biosynthetic route, resulting in such low titers ^2^. A synthetic scaffold was proven effective in increasing MIOX stability and the efficiency of the biosynthetic route, leading to 2.5 g/L D-glucaric acid produced from 10 g/L D-glucose ^7^. Moreover, an N-terminal SUMO fusion to MIOX gave rise to a 75% increase in D-glucaric acid production from *myo*-inositol where up to 4.85 g/L of D-glucaric acid was produced from 10.8 g/L *myo*-inositol in recombinant *E. coli* ^4^. However, *E. coli* seems not an excellent candidate for D-glucaric acid production at high titer because D-glucaric acid concentrations above 5 g/L appears to inhibit its further production by *E. coli* through a pH-mediated effect ^2, 3, 5^.

Thus, Gupta et al.^5^ ported the synthetic D-glucaric acid pathway from *E. coli* to *S. cerevisiae*, another model strain widely used industry that has better acid tolerance ^3^, and found that MIOX4 from *Arabidopsis thaliana* outperformed MIOX from *M. musculus* ^5^. The maximal titer in the *S. cerevisiae* strain containing MIOX4 was 1.6 g/L D-glucaric acid from glucose supplemented with *myo*-inositol ^5^. Chen et al.^3^ used delta-sequence-based integrative expression to increase MIOX4 activity and stability, successfully increasing glucaric acid titer about eight times over that of episomal expression. Combining this strategy with fed-batch fermentation supplemented with 60 mM (10.8 g/L) *myo*-inositol, a titer of 6 g/L (28.6 mM) D-glucaric acid was achieved, which was the highest had been ever reported in *S. cerevisiae* ^3^.

Here we used the same genes and strategy as Chen et al. described ^3^ to construct the biosynthetic route for D-glucaric acid production in a different baker’s yeast strain, *S. cerevisiae* INVSc1. In light of *myo*-inositol availability was rate-limiting in the *S. cerevisiae* strain containing *miox4* gene from *A. thaliana* ^3, 5^, the gene of *opi1* in *S. cerevisiae* INVSc1 was also knocked out accordingly to remove its negative regulation on *myo*-inositol synthesis 3, 8. As a result, a more robust engineered strain of *S. cerevisiae* INVSc1 was obtained, producing the record-breaking titer of D-glucaric acid in *S. cerevisiae*. As the top value-added chemical from biomass, however, there have been no reports on how to produce D-glucaric acid from lignocellulose in the scenario of biorefinery.

As shown in Fig. 1, we applied the engineered strain of *S. cerevisiae* INVSc1 to bioprocesses for D-glucaric acid production from model cellulose and natural lignocelluloses, the same as biofuel production such as bioethanol, including separated hydrolysis and fermentation (SHF), simultaneous saccharification and fermentation (SSF) and consolidated bioprocessing (CBP) ^9, 10, 11, 12, 13, 14^. SHF and SSF were carried out in the context of on-site cellulase production because this mode has many advantages, enabling cost saving cost and tailor made enzyme for a given feedstock ^13, 14, 15^. CBP by an artificial microbial consortium composed of *Trichoderma reesei* Rut-C30 and the engineered *S. cerevisiae* strain was established in this study to actualize direct production of D-glucaric acid from lignocelluloses. Unlike the CBP by the same microbial consortium consisting of *T. reesei* and *S. cerevisiae* where the ethanol production by *S. cerevisiae* required anaerobic condition but *T. reesei* was aerobic ^12^, the CBP here was simpler because both the cellulase production by *T. reesei* and D-glucaric acid biosynthesis by *S. cerevisiae* were aerobic. The artificial microbial consortium for CBP was attempted to be optimized and the team of the two strains was found to be the best with excellent work distribution and collaboration. CBPs by the artificial microbial consortium achieved efficient D-glucaric acid production of lignocelluloses at a near gram per liter level. This work provides an example for production of D-glucaric acid at record-breaking titer and presents the discovery that CBP of lignocellulose by the artificial microbial consortium is a desirable and promising approach for D-glucaric acid production.

**Fig. 1.**
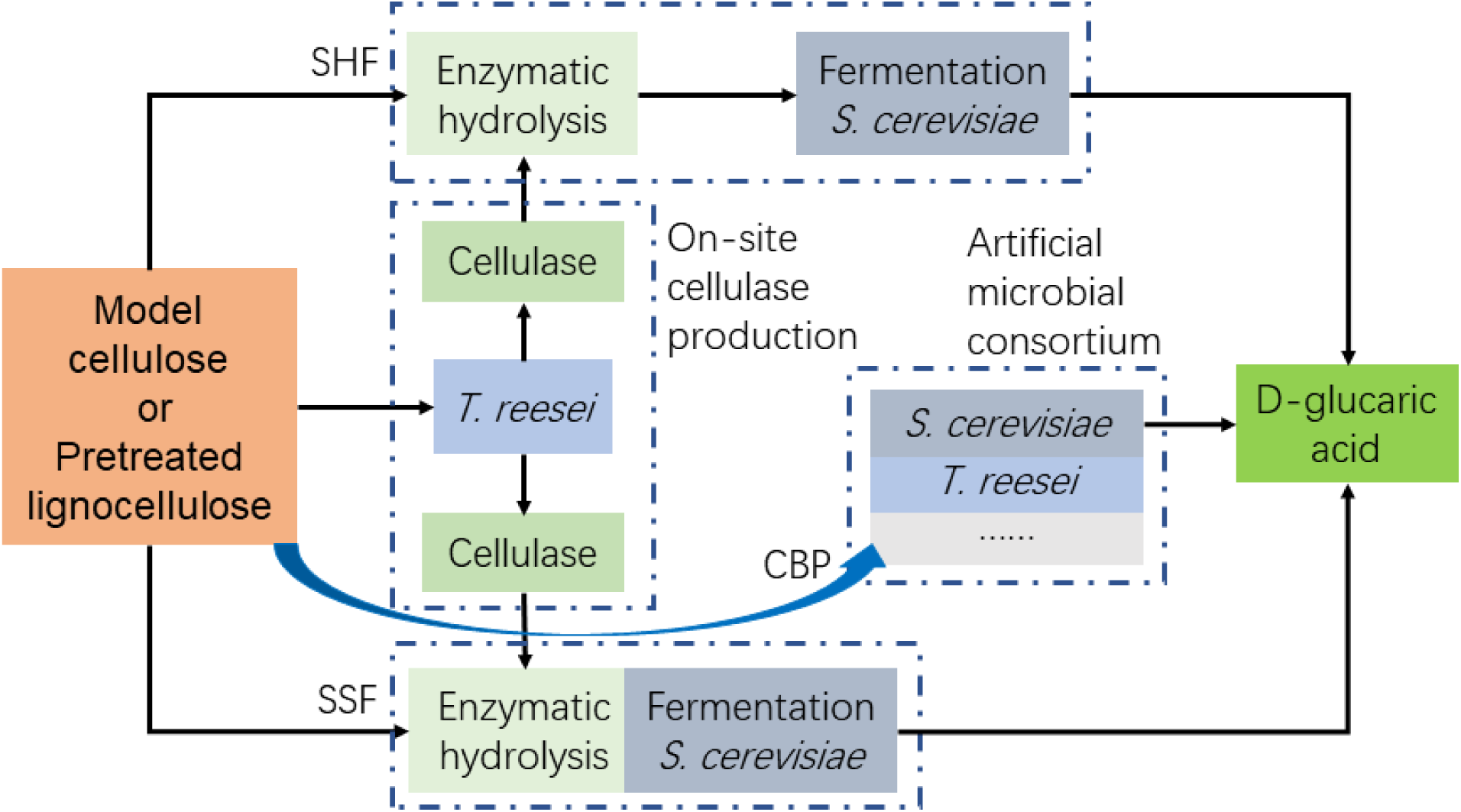
Diagram of biotechnological D-glucaric acid production from lignocellulose. SHF, separated hydrolysis and fermentation; SSF, simultaneous saccharification and fermentation; CBP, consolidated bioprocessing.

## Methods

### Plasmids and their construction

The two heterologous genes, *miox4* gene encoding MIOX4 enzyme in *A. thaliana* ^16^ and the *udh* gene encoding UDH in *P. syringae* ^17^, were codon-optimized according to the codon preference of *S. cerevisiae* and synthesized by Sangon Biotech (Shanghai, China). Then these two genes were ligated to the plasmid of pY26-GPD-TEF, purchased from Miaoling Bioscience & Technology Co., Ltd. (Wuhan, China), using the combinations of restriction enzymes, *Bgl*II/*Not*I and *Eco*RI/*Xho*I, to generate pY26-miox4-udh (Table S1). This recombinant vector carrying *miox4* and *udh* under the control of the promoters *Pgpd* and *Ptef*, respectively, were used for the subsequent transformation of *S. cerevisiae*.

pUG6 from Miaoling Bioscience & Technology Co., Ltd. (Wuhan, China) with kanamycin resistance gene was used as the template for the construction of the knock out cassette loxP-Kan-loxP by polymerase chain reaction (PCR) using the primers knock-OPI1F/R (Table S2). The plasmid pSH47 from Miaoling Bioscience & Technology Co., Ltd. (Wuhan, China) provided Cre recombinase for the self-recombination of the knock out cassette.

The plasmid pCAMBIA1300 ^18, 19^ was used as the backbone for constructing the recombinant vector pCA-Pcbh1-ips-Tcbh1 (Table S1), where the gene *ips* (GenBank: L23520.1) encoding *myo*-inositol-1-phosphate synthase from *S. cerevisiae* ^2^ was codon-optimized according to the codon preference of *T. reesei* and expressed heterologously under the control of the strong promoter *Pcbh1* to improve the *myo*-inositol production.

### Strains and media

All strains used in this work, including the starting strains and the engineered strains, are listed in Table S1.

LB medium, containing 1 g/L tryptone, 0.5 g/L yeast extract, and 1 g/L NaCl, was used to culture *E. coli* cells and *Agrobacterium tumefaciens* cells after being autoclaved at 121□ for 20 min. YPD medium was used to culture *S. cerevisiae* cells, which had a following composition (g/L): 10 yeast extract, 20 peptone, and 20 glucose. When selective YPD medium was prepared, geneticin G418 was added to a specific concentration after being autoclaved and cooled down. Solid media were prepared by adding 2 g/L agar before autoclave.

The seed medium for *T. reesei* strains was composed of 10 g/L glucose, 1 g/L peptone, 5 mL Mandels nutrient salts solution ^20^, 2.5 mL citrate buffer (1 mol/L), 0.05 mL Mandels trace elements solution ^20^, and 0.1 g/L Tween 80. The seed medium was autoclaved at 121□ for 20 min. The fermentation medium for cellulase production by *T. reesei* comprised of 15 g/L Avicel or 30 g/L pretreated lignocellulose (dry biomass), 1 g/L glucose, 6 g/L (NH_4_)_2_SO_4_, 2.0 g/L KH_2_PO_4_, 0.3 g/L CaCl_2_, 0.3 g/L MgSO_4_, 0.005 g/L FeSO_4_, 0.0016 g/L MnSO_4_, 0.0014 g/L ZnSO_4_ and 0.0037 g/L CoCl_2_. The initial pH was adjusted to 4.8 with citrate buffer. This fermentation medium was autoclaved at 121□ for 30 min.

All chemicals except lignocellulosic materials were purchased from Sinopharm Chemical Reagent Co. Ltd., Shanghai, China. The media for SSF and CBP were detailed in the Methods sections about them.

### Genetic engineering of *S. cerevisiae*

The cassette loxP-Kan-loxP amplified from pUG6 by PCR using the primers knock-OPI1F/R (Table S2) was transformed into the competent cells of *S. cerevisiae* INVSc1 prepared with the Li-Ac method ^21^ to knock out *opi1*. The *KanMX* gene disruption cassette was cured by the homologous recombination between the loxP sites mediated by Cre recombinase expressed by the plasmid pSH47. This plasmid pSH47 was lost as the host cells were cultivated continuously, resulting in the *S. cerevisiae* INVSc1 *opi1*Δ strain which was used as the host for constructing the biosynthetic pathway for D-glucaric acid production.

For this target, there were two ways to express foreign genes, episomal or integrative expression. In the scenario of the episomal expression, the recombinant vector pY26-miox4-udh was directly transformed into the competent cells of *S. cerevisiae* INVSc1 *opi1*Δ and the high glucaric acid-producing strain was screened. For the integrative expression, the overlap PCR was conducted to splice the expression cassettes with delta1 and delta2, targeting the integrations into Ty loci ^3, 22^. First, delta1 and delta2 were amplified from *S. cerevisiae* genome using the primers delta1-F/delta1-R and delta2-F/delta2-R (Table S2) respectively, and the *miox4* and *udh* expression cassettes were amplified from the plasmid pY26-miox4-udh using the primers MIXO4-F/R and UDH-F/R respectively. Second, the fragments d1-M-F and L-U-d2 were generated by the overlap PCR from delta1 and MIXO4 expression cassette using the primers delta1-F and FURA3-R and from UDH expression cassette and delta2 using the primers LURA-F/delta2-R, respectively. Finally, the whole fragment was gained from the overlap PCR from the fragments d1-M-F and L-U-d2 using the primers delta1-F and delta2-R, which was purified with Cyle-Pure Kit 200 (Omega Bio-tek, Georgia, USA) and used for the transformation of the competent cells of *S. cerevisiae* INVSc1 *opi1*Δ.

### Genetic engineering of *T. reesei*

The recombinant plasmid pCA-Pcbh1-ips-Tcbh1 was transformed into *T. reesei* Rut-C30 by the method of *A. tumefaciens* mediated transformation (AMT) ^23^. Then the potential *T. reesei* transformants were selected by the two rounds of screening, the first PDA (potato dextrose agar) plates added with hygromycin B and the second Avicel plates as described above ^18, 19^. The fast-growing *T. reesei* transformants selected by the two rounds of screening were tested in the fermentation for *myo*-inositol production and used in the artificial microbial consortium to increase *myo*-inositol availability.

### Fermentation for D-glucaric acid production by the engineered *S. cerevisiae*

Fermentation was carried out in 250 mL Erlenmeyer flasks with a working volume 50 mL of YPD medium with or without 10.8 g/L (60 mM) *myo*-inositol. Before inoculation into fermentation medium, *S. cerevisiae* strains were precultured in 5 mL YPD medium with 10.8 g/L (60 mM) *myo*-inositol in 50 mL Erlenmeyer flasks at 30□ with a shaking of 250 rpm to an optical density at 600 nm (OD_600_) of ∼5. Then, the cells were collected and inoculated into fermentation medium with or without 10.8 g/L (60 mM) *myo*-inositol to an OD_600_ of 0.1. Fermentation was implemented at 30□ with a shaking of 250 rpm.

Fed-batch fermentation was conducted under the same conditions as the batch fermentation mentioned above, except that 5 g/L glucose was supplemented twice during the fermentation process, 24 h and 48 h respectively.

### Separated hydrolysis and fermentation (SHF)

In the scenario of SHF (Fig. 1), the cellulase was produced by *T. reesei* Rut-C30 from Avicel or steam exploded corn stover (SECS) ^14, 24, 25^ was applied to the enzymatic hydrolysis of themselves, so called “on-site cellulase production” ^13, 14, 15, 25^. *T. reesei* Rut-C30 was precultured in the seed medium for 36 h and then the seed collected and inoculated into the fermentation medium for cellulase production. The fermentation broth containing crude cellulase was directly used in the enzymatic hydrolysis of Avicel or SECS ^14, 19^, which was operated in 250 mL Erlenmeyer flasks with a working volume of 50 mL 2.5 mL 1 M citrate buffer solution (for final pH 4.8), 50 g/L Avicel or 100 g/L SECS (dry material), 25 FPIU/g glucan the cellulase harvested after 5 d fermentation, and a supplementary amount of water to make up 50 mL. The enzymatic hydrolysis was conducted in a rotary shaker at 50□ with a shaking of 140 rpm for 48 h. The resulted enzymatic hydrolysates containing fermentable sugars were used to comprise of fermentation medium supplemented with the same nutrients as YPD medium except glucose. The following operations were the same as those described in the section of fermentation for D-glucaric acid production by the engineered *S. cerevisiae*.

### Simultaneous saccharification and fermentation (SSF)

In the scenario of SSF (Fig. 1), the cellulase prepared by the same method in SHF was used in the enzymatic prehydrolysis of Avicel or SECS for 12 h. The prehydrolysis of SSF was performed in 250 mL Erlenmeyer flasks with 50 mL SSF reaction mixture containing 50 g/L Avicel or 100 g/L SECS (dry material), 25 FPIU/g glucan the cellulase harvested after 5 d fermentation, 6 g/L (NH_4_)_2_SO_4_, 2.0 g/L KH_2_PO_4_, 0.3 g/L MgSO_4_·7H_2_O, 0.3 g/L CaCl_2_·2H_2_O, 0.1 g/L Tween 80, 10 g/L peptone, 5 g/L yeast extract,, and a supplementary amount of water to make up 50 mL. The initial pH of the reaction mixture was adjusted to 4.8 with citrate buffer. The reaction mixture without cellulase was autoclaved at 121□ for 30 min. After cellulase addition, the reaction mixture was incubated at 50□ with a shaking of 140 rpm for 12 h. Then, the temperature was decreased to 33□ (or specified in the main text when studying its effect on SSF) and the shaking was increased to 250 rpm. The precultured *S. cerevisiae* with an OD_600_ of ∼5 was inoculated into the reaction mixture for the SSF to produce D-glucaric acid.

### Consolidated Bioprocessing (CBP)

The medium for CBP had a following composition: 15 g/L (or specified in the main text when investigating its effect on CBP) Avicel or SECS (dry material), 1 g/L peptone, 1 g/L yeast extract, 10%(v/v) Mandels nutrient salts solution ^20^, 0.1%(v/v) trace elements solution ^20^, 5%(v/v) citrate buffer (1 mol/L), 0.1 g/L Tween 80. CBP medium was autoclaved at 121□ for 30 min. *T. reesei* was precultured in the seed medium at 30□ for 36 h and *S. cerevisiae* was precultured in YPD medium at 30□ till an OD_600_ of ∼5. Then *T. reesei* was inoculated into CBP medium and the inoculation of *S. cerevisiae* was implemented immediately or delayed (or specified in the main text when studying its effect on CBP). The inoculum ratio of *T. reesei* to *S. cerevisiae* was 1:1 (or specified in the main text when studying its effect on CBP). If other strain or species was inoculated, the detailed information would be specified in the main text. The total inocula were 10%(v/v) of the fermentation medium. CBP was carried out in 250 mL Erlenmeyer flasks with 50 mL medium at 30□ with a shaking of 180 rpm. *A. niger* was precultured for 48 h by the same method as *T. reesei* if needed by CBP.

### Analytical methods

Filter paper activity (FPA) of cellulase, representing the total enzymatic activity, was assayed by the method standardized by the International Union of Pure and Applied Chemistry (IUPAC) ^26^, which quantifies the total amount of the reducing sugars produced from 50 mg Whatman No.1 filter paper (1×6 cm strip) by cellulase within 60 min. One International Unit of FPA (FPIU) was defined as the amount of cellulase needed for producing 1 μmol reducing sugars in 1 min.

β-Glucosidase activity (BGA) was determined using the standard method ^26^ with the tiny modification, i.e. the substrate ρNPG (ρ-nitrophenyl-β-d-1,4-glucopiranoside) (Sigma-Aldrich, St. Louis, MO, USA). The amount of ρ-nitrophenol produced from ρNPG by β-glucosidase within 10 min was assayed using spectrophotometer at a wavelength of 400 nm. One International Unit of BGA (IU) was defined as the amount of β-glucosidase required for producing 1 μmol of ρ-nitrophenol from ρNPG in 1 min.

Cellobiohydrolase activity (BGA) was assayed according to the method modified from FPA measurement method ^26^. Microcrystalline cellulose PH101 purchased from Sigma-Aldrich (St. Louis, MO, USA) in the form of 1% (w/v) suspension was used as the substrate for the reaction with the duration of 30 min. One Unit (1 U) of CBA was defined as the amount of enzyme required for producing 1 mg reducing sugars in 1 h.

High performance liquid chromatography (HPLC) was adopted to analyze and quantify D-glucaric acid, *myo*-inositol and sugars, where Shimadzu Prominence LC-20A system equipped with Bio-Rad Aminex HPX-87H (300 mm×7.8 mm) column was used. When D-glucaric acid was determined, an ultraviolet (UV) detector was employed to detect the eluate. While *myo*-inositol and sugars were determined, a refractive index (RI) detector was employed. Sulfuric acid (5 mmol/L) was used as the mobile phase and set at a flow rate of 0.6 mL/min. The column temperature was maintained at 50□. Sample loading for each injection was 10 μL.

Liquid Chromatography-Mass Spectrometry (LC-MS) was used to characterize and identify D-glucaric acid, where Shimadzu L-30A and AB Sciex Triple TOF 5600 were employed. The column was Shimadzu Shim-pack XR-ODS 100L×2.0. The mobile phases, set at a flow rate of 0.15 mL/min, were the aqueous solution containing 1 mmol/L ammonium formate and 1%(v/v) formic acid (Mobile phase A) and the acetonitrile solution containing 1 mmol/L ammonium formate (Mobile phase B), respectively. Mobile phase B, B for short and the same below, was used as the eluent and the elution procedure was as follows: 0-12 min 30% B, 12-30 min 30%-65% B, 30-31 min 65%-95% B, 31-35 min 95% B, 35-36 min 95%-30% B, 36-40 min 30% B. Sample loading for each injection was 5 μL. Negative electrospray ionization mode was chosen to ionize samples. The flow rate of atomizing gas (N_2_) was 1.5 L/min. The temperature for CDL and HB was 200□. Scanning scope (m/z) ranged from 150 to 300.

## Results

### Construction of D-glucaric acid-producing *S. cerevisiae*

In order to relieve the negative regulation of *opi1* on *myo*-inositol synthesis, *opi1* was knocked out to generate *S. cerevisiae* strain INVSc1 *opi*Δ, which was subsequently used as the starting strain for the construction of D-glucaric acid-producing *S. cerevisiae*, as detailed in Methods. The biosynthetic pathway of D-glucaric acid in the engineered *S. cerevisiae* is illustrated in Fig. 2A. Two D-glucaric acid-producing strains, *S. cerevisiae* LGA-1 with the integrative expression of foreign genes and *S. cerevisiae* LGA-C with the episomal expression, were chosen for subsequent fermentation experiments. The products of *S. cerevisiae* LGA-1 and LGA-C were characterized by LC-MS to verify their capability of producing D-glucaric acid and the results are confirmative, as shown in Fig. S1.

**Fig. 2.**
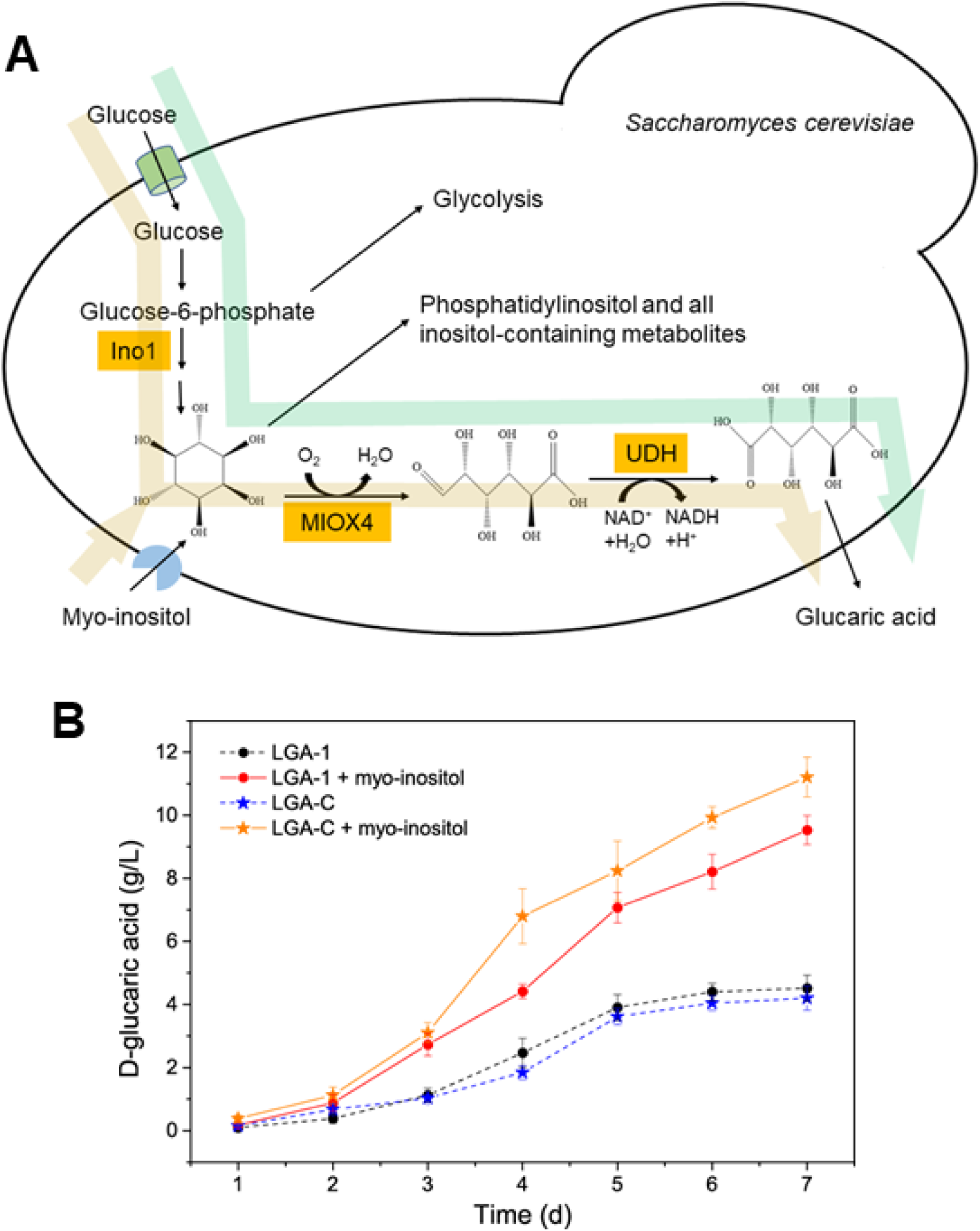
(A) Biosynthetic pathway of D-glucaric acid introduced into *S. cerevisiae*. Ino1, *myo*-inositol-1-phosphate synthase; MIOX4, *myo*-inositol oxygenase; UDH, urinate dehydrogenase. Green arrow illustrates D-glucaric acid production from glucose, and yellow one shows D-glucaric acid production from glucose and *myo*-inositol. (B) Time courses of fed-batch fermentations on YPD medium without and with 10.8 g/L *myo*-inositol. LGA-1 and LGA-C are the engineered *S. cerevisiae* strains capable of producing D-glucaric acid. Data shown here are average values of at least three biological replicates and error bars are standard deviations.

### Fed-batch fermentation with or without *myo*-inositol

These two strains, *S. cerevisiae* LGA-1 and *S. cerevisiae* LGA-C, were compared in the fermentations on YPD medium without and with 10.8 g/L (60 mM) *myo*-inositol which was reported to have the highest D-glucaric acid titer ^3, 5^. The time courses of the fermentations are shown in Fig. 2B. It was found that the fermentation processes supplemented with *myo*-inositol had higher D-glucaric acid titer than those using glucose as sole carbon source. This observation is the same as the previous reports ^3, 5^. LGA-1 performed similarly to LGA-C when using glucose as sole carbon source, in contrast to their performances in the fermentation supplemented with *myo*-inositol where LGA-C outperformed LGA-1. After 7 d fermentation, LGA-1 produced 9.53 ± 0.46 g/L and LGA-C produced 11.21 ± 0.63 g/L D-glucaric acid from 30 g/L glucose and 10.8 g/L *myo*-inositol in the mode of fed-batch fermentation, respectively.

Production of D-glucaric acid from *myo*-inositol (marked with yellow arrows in Fig. 2A), however, is not a desirable approach because the latter is also an industrially value-added chemical ^27, 28^. Moreover, so far there has not been relevant reports pertaining to D-glucaric acid production from lignocellulose, though it was considered as one of the top value-added chemicals from biomass ^1^. Therefore, production of D-glucaric acid from lignocellulose in the context of biorefinery, the same as production of lignocellulosic biofuels such as lignocellulosic ethanol ^9, 10^, should be treated as an important attempt to seek cheaper manufacturing of D-glucaric acid. Hence, LGA-C and LGA-1 were applied to the D-glucaric acid production from Avicel and pretreated corn stover via SHF, SSF and CBP, typical bioprocesses in biorefinery.

### Separated hydrolysis and fermentation (SHF)

The time courses of cellulase production from Avicel and SECS are presented in Fig. 3A. Avicel induced higher FPA than SECS in the cellulase production by *T. reesei* Rut-C30. On Day 5, the FPAs induced by Avicel and SECS were 2.87 ± 0.29 and 2.45 ± 0.36 FPIU/mL, respectively. Then FPAs decreased because *T. reesei* Rut-C30 entered decline phase, which is in line with the previous work ^18, 29^. Thus, the cellulases harvested on Day 5 were applied to the enzymatic hydrolysis of themselves in the context of on-site cellulase production ^13, 14, 15^, i.e. the cellulase induced by Avicel used for the enzymatic hydrolysis of Avicel, because this is advantageous over the cellulases induced by other substrates or commercial cellulases ^15, 30^. The results of the enzymatic saccharification of Avicel and SECS are shown in Fig. 3B. Enzymatic hydrolysis of SECS was found to have higher yield than that of Avicel, the former producing 39.73 ± 0.95 g/L glucose from 100 g/L SECS containing 53.2 g/L glucan and the latter 28.31 ± 1.17 g/L glucose from 50 g/L Avicel. The resulted hydrolysates containing fermentable sugars were subsequently fermented by the engineered *S. cerevisiae* strains, LGA-1 and LGA-C, for D-glucaric acid production. The time courses are shown in Fig. 3C, which were in the similar pattern. Though the enzymatic hydrolysate of SECS fermented by LGA-1 led to the highest D-glucaric acid titer of 4.92 ± 0.24 g/L after 7 d fermentation, the differences were not so distinguishable, let alone the enzymatic hydrolysate of Avicel had lower concentration of glucose.

**Fig. 3.**
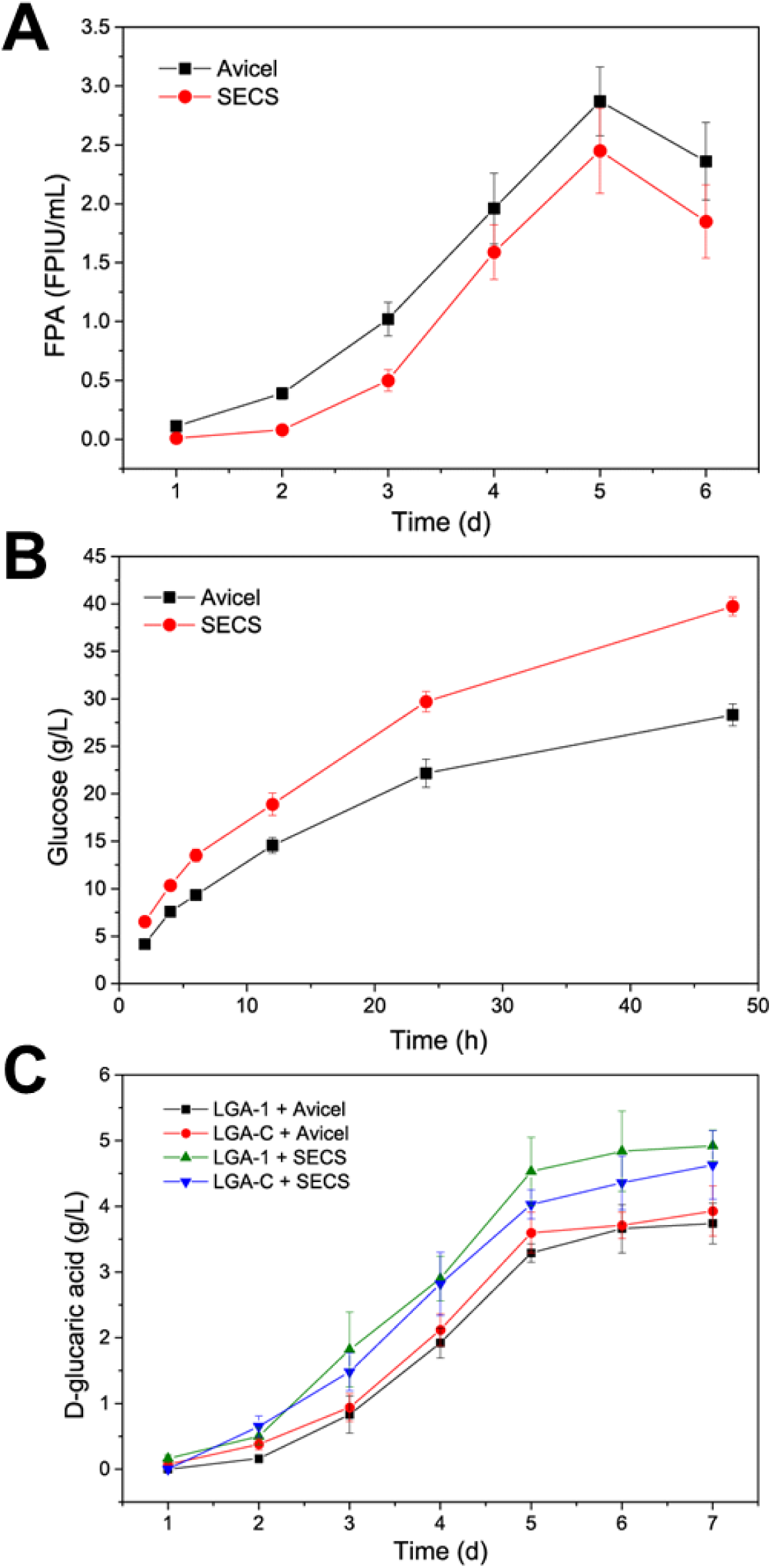
(A) Time courses of cellulase production by *T. reesei* Rut-C30 from Avicel and steam-exploded corn stover (SECS). (B) Enzymatic hydrolysis of Avicel and SECS by the cellulases produced in the context of on-site cellulase production. (C) Time courses of batch fermentations on Avicel hydrolysate and SECS hydrolysate. LGA-1 and LGA-C are the engineered *S. cerevisiae* strains capable of producing D-glucaric acid. Data shown here are average values of at least three biological replicates and error bars are standard deviations.

Overall, LGA-1 produced 3.74 ± 0.31 g/L D-glucaric acid and LGA-C produced 3.93 ± 0.38 g/L D-glucaric acid from 50 g/L Avicel. LGA-1 produced 4.92 ± 0.24 g/L D-glucaric acid and LGA-C produced 4.63 ± 0.52 g/L D-glucaric acid from 100 g/L SECS.

### Simultaneous saccharification and fermentation (SSF)

Three temperatures, 30, 33 and 36□, were tested here to study the effect on SSF and the results are shown in Fig. S2, Fig. 4 and Fig. S3, respectively. It was found that 33□ was the most suitable temperature for SSF, leading to the highest concentrations of D-glucaric acid. This is the same as SSF for bioethanol production where 33□ was also the most suitable temperature ^11^.

**Fig. 4.**
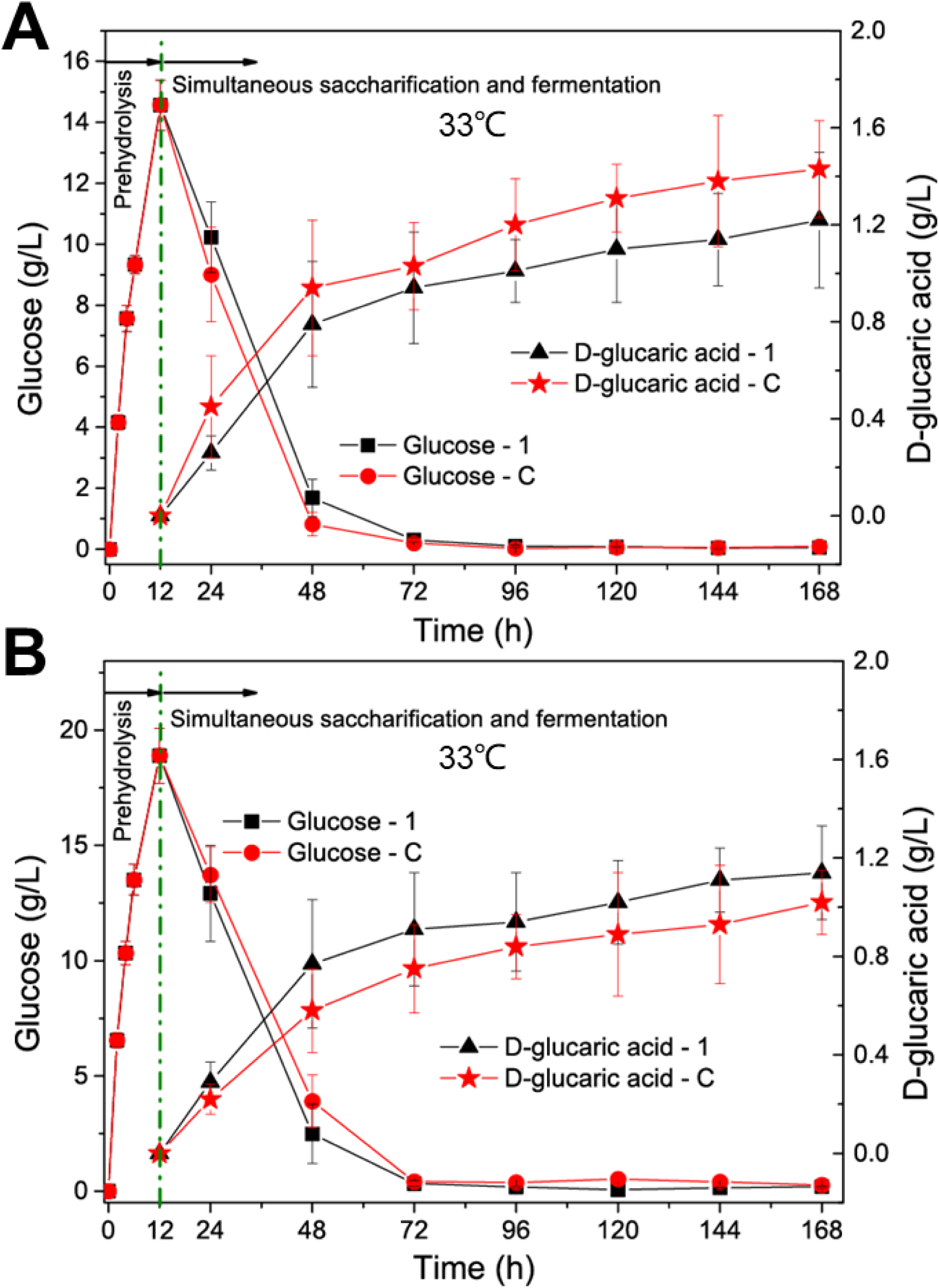
Simultaneous saccharification and fermentation of Avicel (A) and steam-exploded corn stover (SECS) (B) at 33□ from 12 h to 168 h after enzymatic prehydrolysis at 50□ for 12 h. LGA-1 and LGA-C are the engineered *S. cerevisiae* strains capable of producing D-glucaric acid. Data shown here are average values of at least three biological replicates and error bars are standard deviations.

The time courses of SSF for D-glucaric acid production from Avicel and SECS are shown Fig. 4A and B, respectively, which displayed similar pattern that glucose dropped rapidly after LGA-1 or LGA-C being inoculated. After 48 h, glucose reduced to an extremely low level close to zero. The titers of D-glucaric acid increased quickly after 12 h, the time point of *S. cerevisiae* inoculation, and reached the plateau after 72 h. LGA-1 and LGA-C had reverse results in the SSF for producing D-glucaric acid, the former produced the higher titer of D-glucaric acid than the latter from Avicel but the former produced lower titer from SECS. However, the differences were not so enormous.

After 7 d, LGA-1 produced 1.22 ± 0.28 g/L D-glucaric acid and LGA-C produced 1.43 ± 0.20 g/L D-glucaric acid from 50 g/L Avicel. LGA-1 produced 1.14 ± 0.19 g/L D-glucaric acid and LGA-C produced 1.02 ± 0.13 g/L D-glucaric acid from 100 g/L SECS.

### Consolidated bioprocessing (CBP)

CBP is a desirable approach for D-glucaric acid production from lignocellulose because both cellulase production by *T. reesei* and D-glucaric acid production by *S. cerevisiae* are aerobic, unlike CBP for bioethanol production where an anaerobic condition should be created for *S. cerevisiae* ^12^. Here the operation of CBP was much easier. CBP for D-glucaric acid production from lignocellulose by *T. reesei* and *S. cerevisiae* is illustrated in Fig. 5A. The microbial consortium composed of *T. reesei* Rut-C30 and the engineered *S. cerevisiae* could achieve the bioconversion of lignocellulose to D-glucaric acid in one step.

**Fig. 5.**
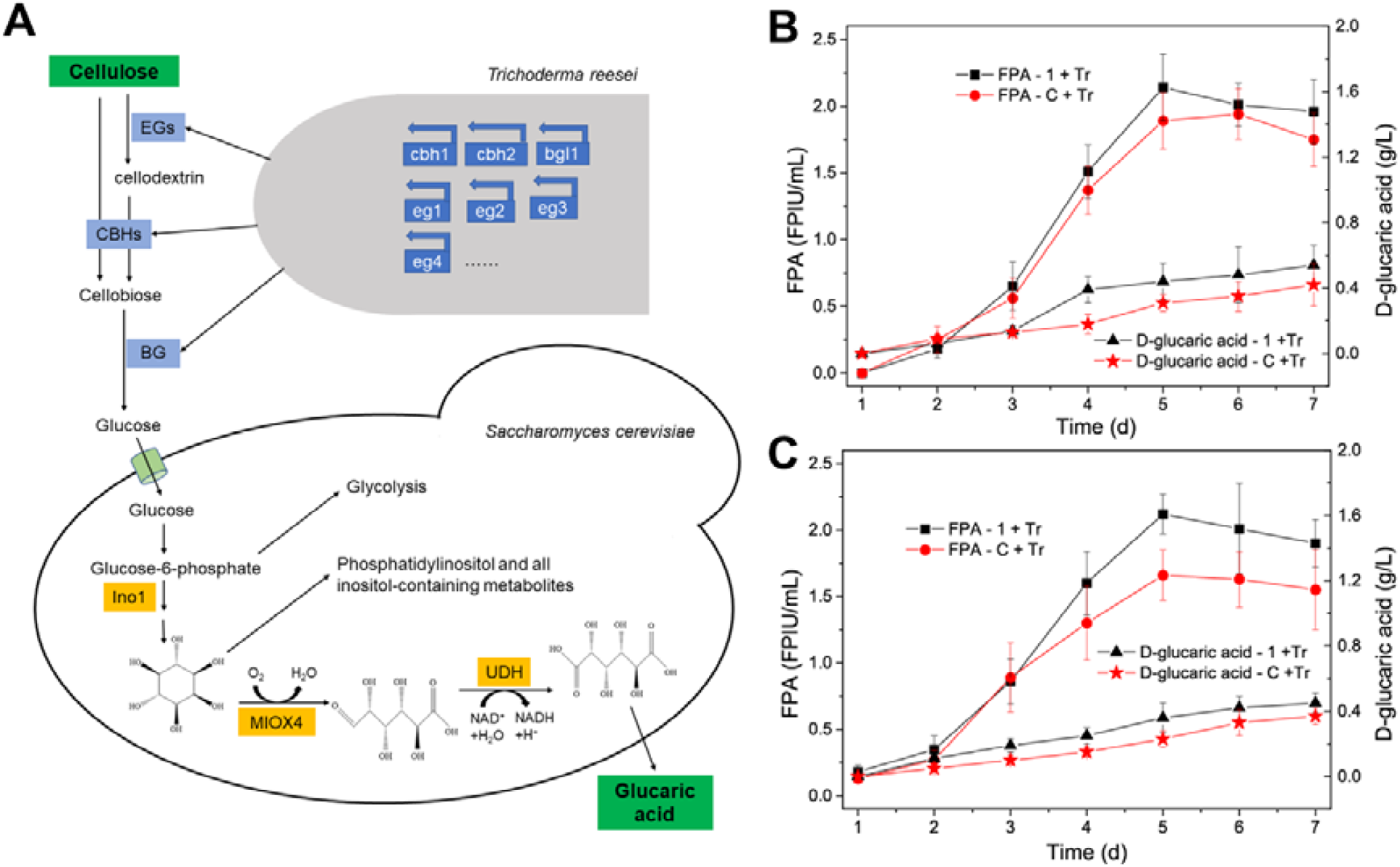
(A) Diagram of CBP of cellulose for D-glucaric acid production by the microbial consortium consisted of *T. reesei* Rut-C30 and *S. cerevisiae* LGA-1. EGs, endoglucanases; CBHs, cellobiohydrolases; BG, β-glucosidase; Ino1, *myo*-inositol-1-phosphate synthase; MIOX4, *myo*-inositol oxygenase; UDH, uronate dehydrogenase. (B) FPAs and concentrations of D-glucaric acid during CBP of Avicel. (C) FPAs and concentrations of D-glucaric acid during CBP of steam-exploded corn stover (SECS). 1 and C in legends stand for *S. cerevisiae* LGA-1 and LGA-C, respectively. Tr in legends stands for *T. reesei* Rut-C30. Data shown here are average values of at least three biological replicates and error bars are standard deviations.

The time courses of CBPs from Avicel and SECS are shown in Fig. 5B and C, respectively. It was found that the microbial consortium of *T. reesei* Rut-C30 and *S. cerevisiae* LGA-1 produced 0.54 ± 0.12 g/L D-glucaric acid from 15 g/L Avicel, and 0.45 ± 0.06 g/L D-glucaric acid from 15 g/L SECS after 7 d fermentation. Somewhat lower concentrations of D-glucaric acid were produced when *S. cerevisiae* LGA-C was used. Thus, *S. cerevisiae* LGA-1 was more desirable in the CBP for D-glucaric acid production. Increase in substrate loading did not lead to higher concentrations of D-glucaric acid, as shown in Fig. S4. The optimal substrate loading in CBP for D-glucaric acid here was lower than that in CBP for bioethanol, 17.5 g/L ^12^. This result implicates the low efficiency of the microbial consortium comprising of *T. reesei* Rut-C30 and *S. cerevisiae* LGA-1 in CBP of lignocellulose to D-glucaric acid.

Further work was thus carried out to improve the efficiency. The effect of the ratio of *T. reesei* to *S. cerevisiae* on CBP was investigated and the results are shown in Fig. S5. Among the inoculum ratios of *T. reesei* to *S. cerevisiae*, 1:1, 1:3, 1:5, 5:1 and 3:1, 1:1 was found to be the best for CBP. Changing the ratio caused lower D-glucaric acid titer, proven incapable of improving the efficiency. The effect of the delay time of *S. cerevisiae* inoculation on CBP was also studied and the results are presented in Fig. S6. Among the delay times of *S. cerevisiae* inoculation, 0, 24 and 48 h, 0 (i.e. *T. reesei* and *S. cerevisiae* were inoculated simultaneously) was found to be the most suitable.

In the light of the mixed culture of *T. reesei* and *A. niger* for enhanced cellulase production ^29, 31^, *A. niger* was introduced into the microbial consortium to improve the efficiency of cellulose degradation. The ratio of *T. reesei* to *A. niger* to *S. cerevisiae* was 5:1:5 and the ratio of the total inocula to fermentation medium was 10%(v/v). The results are presented in Fig. 6A, D, E, F and G, and it was found that the microbial consortium composed of *T. reesei, A. niger* and *S. cerevisiae* produced lower concentrations of D-glucaric acid than the microbial consortium consisted of *T. reesei* and *A. niger*, though the former produced higher FPA. This suggests that the microbial consortium of *T. reesei, A. niger* and *S. cerevisiae* was not suitable for CBP of lignocellulose for D-glucaric acid production.

**Fig. 6.**
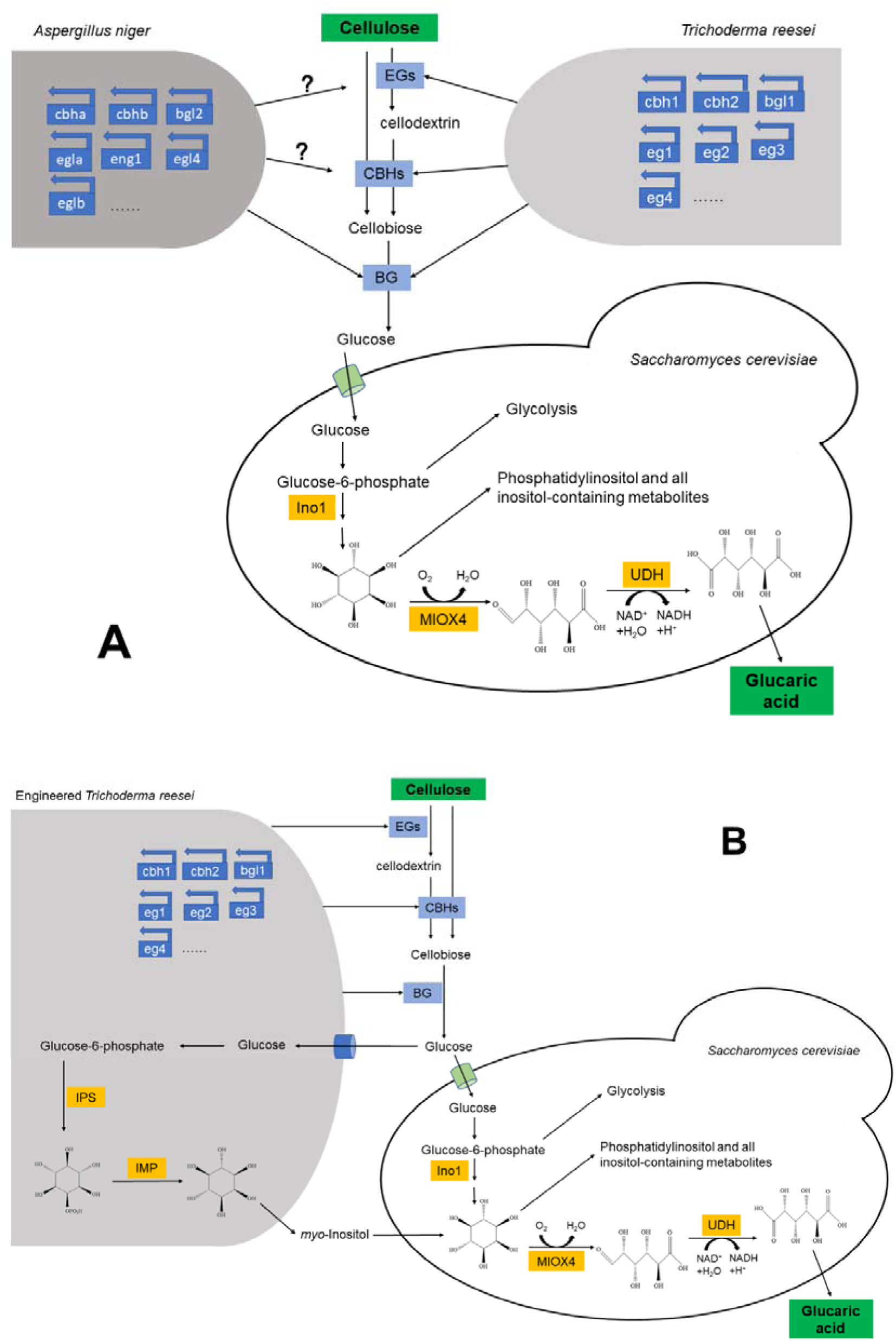
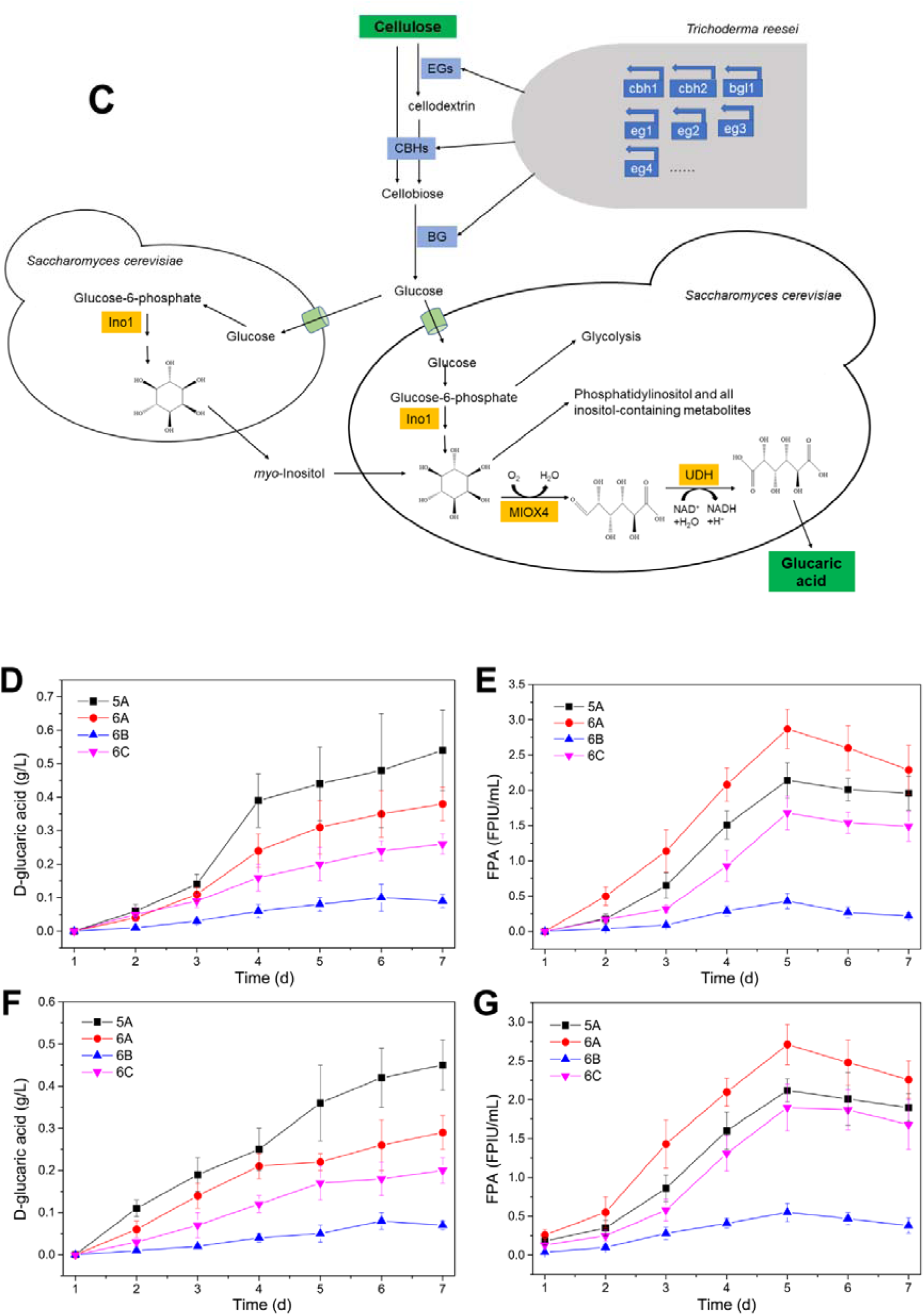
(A) Diagram of CBP of cellulose for D-glucaric acid production by the microbial consortium consisted of *T. reesei* Rut-C30, *A. niger* CICC2103 and *S. cerevisiae* LGA-1. (B) Diagram of CBP of cellulose for D-glucaric acid production by the microbial consortium consisted of the engineered *T. reesei* and *S. cerevisiae* LGA-1. (C) Diagram of CBP of cellulose for D-glucaric acid production by the microbial consortium consisted of *T. reesei* Rut-C30, *S. cerevisiae* Δ*opi1* and *S. cerevisiae* LGA-1. Concentrations of D-glucaric acid (D) and FPAs (E) during CBP of Avicel. Concentrations of D-glucaric acid (F) and FPAs (G) during CBP of steam-exploded corn stover (SECS). 5A, 6A, 6B and 6C in the legends of Fig. 6D, E, F and G represents the CBPs illustrated in Fig. 5A, 6A, 6B and 6C, respectively. Data shown here are average values of at least three biological replicates and error bars are standard deviations.

Gupta et al ^5^. found that *myo*-inositol availability was the rate-limiting step in the engineered *S. cerevisiae* expressing *miox4* gene from *A. thaliana*. The same strategy was used here. Thus, we adopted the following strategies to increase *myo*-inositol availability. We engineered *T. reesei* Rut-C30 to enable it accumulating extracellular *myo*-inositol by expressing inositol-3-phosphate synthase (*ips*, GenBank: L23520.1) and inositol monophosphatase (*imp*, GenBank: CP029160.1) from *S. cerevisiae*. But this was proven unsuccessful because engineering *T. reesei* to produce *myo*-inositol made it no longer potent in cellulase production, let alone the engineered *T. reesei* produced a negligible concentration of myo-inositol (Fig. S7). The results in Fig. 6B, D, E, F and G indicate that this was the most unsuitable strategy to improve D-glucaric acid production by CBP of lignocellulose.

Then, we turned eyes to the engineered *S. cerevisiae* with *opi1* gene being knocked out, whose myo-inositol accumulation was improved (Fig. S8), i.e. the parental strain for LGA-1 and LGA-C. *S. cerevisiae* Δ*opi1* was added to the microbial consortium to try to improve the efficiency of CBP for D-glucaric acid production. The inoculum ratio of *T. reesei* to *S. cerevisiae* Δ*opi1* to *S. cerevisiae* LGA-1 was 2:1:1 and the total volume of the inocula was 10%(v/v) of fermentation medium. It was found in Fig. 6C, D, E, F and G that this strategy was inferior to that strategy described in Fig. 5A.

Overall, the microbial consortium of *T. reesei* Rut-C30 and *S. cerevisiae* LGA-1 was the most efficient CBP for D-glucaric acid production from lignocellulose. Addition of the third strain or metabolic engineering of *T. reesei* for enhanced *myo*-inositol availability was proven failed in improving the efficiency of CBP.

### Comparison of different biorefinery processes for D-glucaric acid production

The parameters of different biorefinery processes for D-glucaric production from Avicel and SECS, including SHF, SSF and CBP, are listed in Table 1 and 2, respectively. The fermentation processes from pure glucose with or without *myo*-inositol supplementation were used as the references for comparison. It is obvious that fermentation on glucose supplemented with *myo*-inositol had the highest D-glucaric acid titer. Next in line is the D-glucaric acid production from glucose or from the hydrolysates of SECS and Avicel containing glucose. SHF had comparable D-glucaric acid titers with the fermentation from pure glucose without *myo*-inositol supplementation. SSF had lower D-glucaric acid titers from Avicel and SECS than those of SHF, and much lower yields than those of SHF. This suggests that SSF was not a desirable approach for D-glucaric acid production from lignocellulose due to its low efficiency.

**Table 1.**
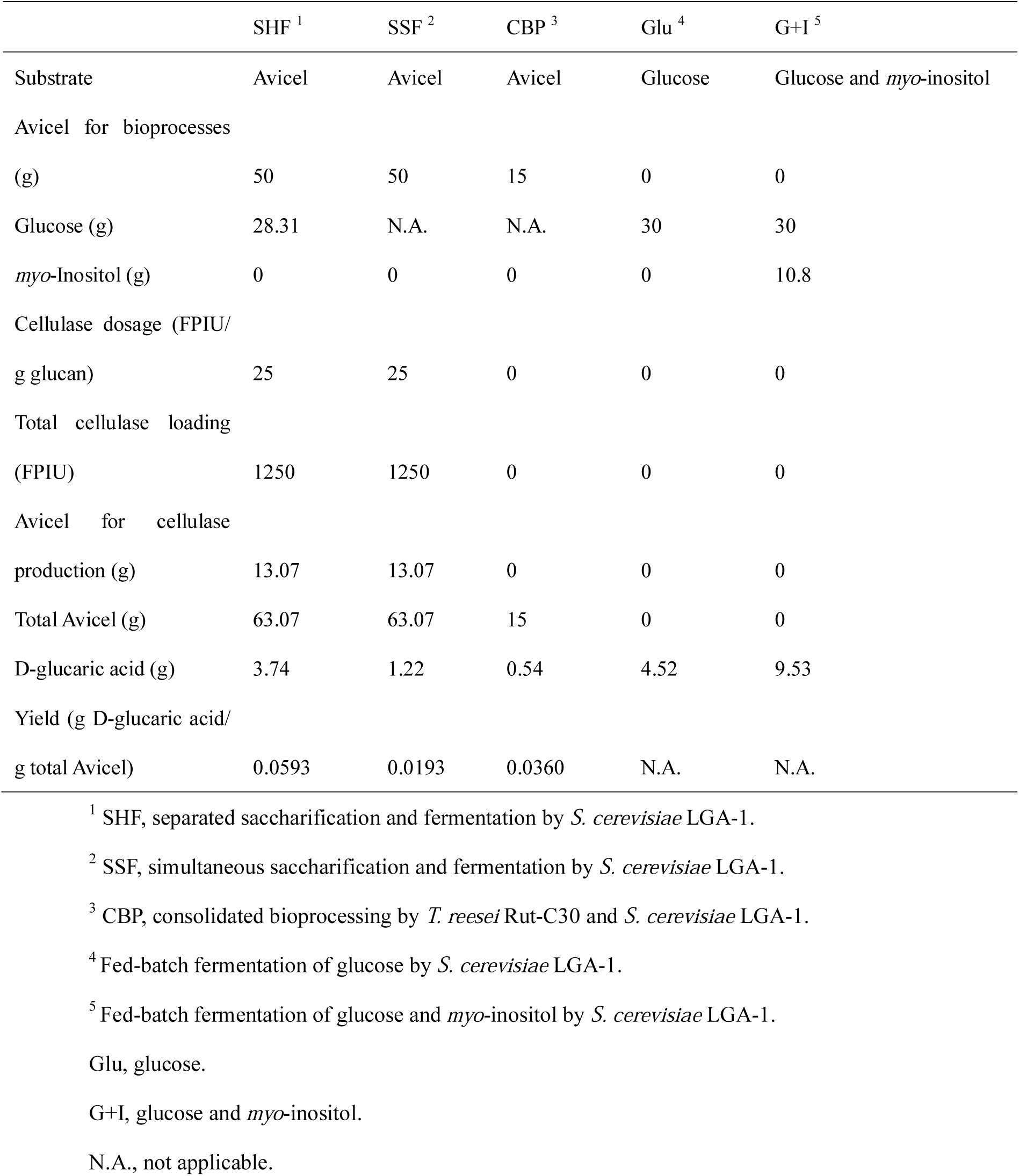
Details on the bioprocesses from Avicel to D-glucaric acid.

**Table 2.**
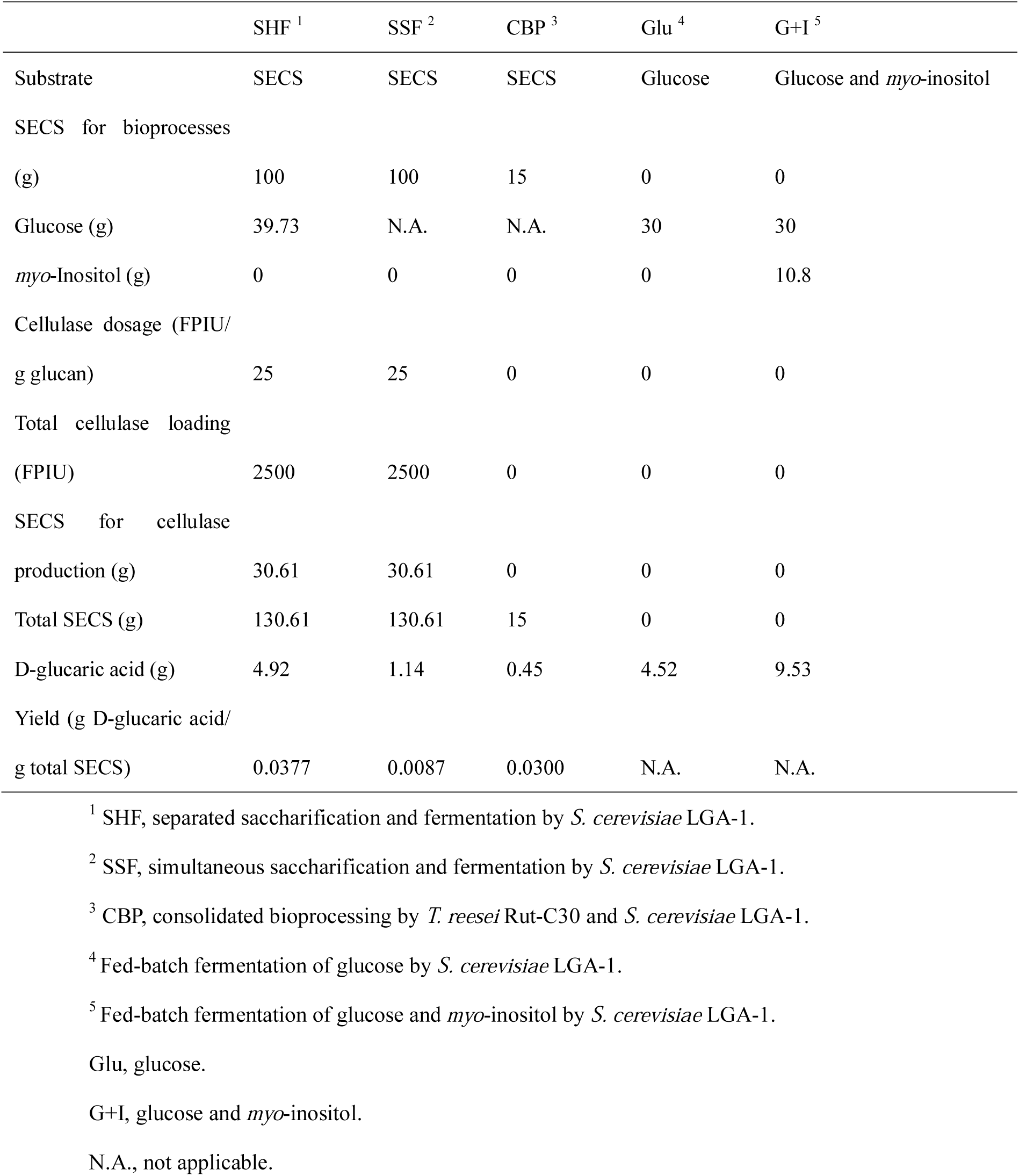
Details on the bioprocesses from steam-exploded corn stover (SECS) to D-glucaric acid.

Though CBP had relatively low D-glucaric acid titer, it had considerably high yields from Avicel and SECS. This is because low substrate loading was used. The yields of CBPs were 0.0360 g D-glucaric acid/g Avicel and 0.0300 g D-glucaric acid/g SECS, much higher than those of SSF and comparable to those of SHF (0.0593 g D-glucaric acid/g Avicel and 0.0377 g D-glucaric acid/g SECS). Moreover, CBP has the least single unit operations and the greatest potential of cost reduction ^10, 12, 32^. In addition, both the cellulase production by *T. reesei* Rut-C30 and D-glucaric acid production by *S. cerevisiae* LGA-1 during CBP of lignocellulose by the microbial consortium are aerobic. This reduces the difficulty and complexity of constructing and operating the microbial consortium ^12^. Therefore, this is highly promising approach for D-glucaric production from lignocellulose.

## Discussion

The strategy of constructing D-glucaric acid-producing *S. cerevisiae* strain with high titer was applied to a different yeast strain *S. cerevisiae* INVSc1 in this work ^3^. Two engineered strains with high titers of D-glucaric acid, LGA-1 and LGA-C, were obtained. LGA-C with the episomal expression of *miox4* and *udh* had higher concentration of D-glucaric acid than LGA-1 with the integrated expression when supplementing *myo*-inositol (Fig. 2B). But no obvious difference observed when using glucose as sole carbon source. This may be because episomal expression was more efficient in transforming *myo*-inositol into D-glucaric acid than integrated expression when abundant *myo*-inositol was available, or the recombinant plasmid with high copy number was led to more MIOX4 and UDH than the integration into Ty loci. In practice, both LGA-1 and LGA-C acted as whole cell catalysts when the fermentation supplemented with *myo*-inositol. In the fed-batch fermentation supplemented with *myo*-inositol, both LGA-1 and LGA-C produced the record-breaking titer of D-glucaric acid, 9.53 ± 0.46 g/L and LGA-C produced 11.21 ± 0.63 g/L D-glucaric acid, much higher than the previous reports on *S. cerevisiae* ^3, 5^ and *E. coli* ^2, 4, 7, 33^. It is noteworthy that higher D-glucaric acid was produced, though we adopted the same strategy as Chen et al ^3^. This is probably because different *S. cerevisiae* strain was used. Different strain means different physiological statuses, different efficiencies of foreign gene expressions, different *myo*-inositol availabilities, etc.

The high titer of D-glucaric acid produced by *S. cerevisiae* LGA-C demonstrates the great potential of *S. cerevisiae* INVSc1 in D-glucaric acid production, although episomal expression is problematic because of its genetic instability ^3^. The higher titer of D-glucaric acid produced from glucose and *myo*-inositol than that from glucose suggests that *myo*-inositol availability is still the rate-liming step for D-glucaric acid production in the engineered *S. cerevisiae*. Furthermore, *myo*-inositol itself is a valuable chemical in industry ^27, 28^. The production of D-glucaric acid from *myo*-inositol was not so economically competitive as the direct production from glucose. In order to enhance the D-glucaric acid production from glucose, therefore, further work should be done to improve the biosynthetic pathway efficiency of D-glucaric acid in *S. cerevisiae*.

As one of the top value-added chemicals from biomass ^1^, there had been no researches on D-glucaric acid production from lignocellulose, the most abundant renewable on this planet. Three biorefinery processes, SHF, SSF and CBP (Fig. 1), were conducted using the engineered *S. cerevisiae* strains, LGA-1 and LGA-C. Two substrates were used in these biorefinery processes, Avicel and SECS. The former is pure cellulose that is often used as model substrate, and the latter is corn stover that is abundant in China but always improperly treated, causing serious air pollution ^30^. The results of SHF (Fig. 3) shows that Avicel induced more cellulase but had lower glucose concentrations. Overall, both D-glucaric acid titer and yield of SECS were comparable to those of Avicel (Table 1 and Table 2). Moreover, the results of SHF were close to fermentation from pure glucose. These indicate that corn stover a suitable feedstock for D-glucaric acid production and the engineered *S. cerevisiae* could be applied to the biorefinery of lignocellulose for D-glucaric production.

SSF was proven not so successful here, leading to the lowest D-glucaric acid titer and yields from Avicel and SECS (Table 1 and 2). This is because SSF is not authentic, which encompassed two phases, enzymatic prehydrolysis and SSF. Only the former phase had the highest hydrolysis efficiency. After entering SSF, temperature was decreased to the appropriate range of *S. cerevisiae* in which the enzymatic hydrolysis rate was reduced. If SSF temperature was increased for improved enzymatic hydrolysis, the fermentation by *S. cerevisiae* was weakened (Fig. 4, Fig. S2 and Fig. S3). The hurdle of SSF is the gap between the optimal temperature for enzymatic hydrolysis by *T. reesei* cellulase and that for fermentation by *S. cerevisiae*. This is the same as SSF for bioethanol production ^11^.

Enlightened by the previous research on CBP for bioethanol production ^12^, we constructed the same microbial consortium of *T. reesei* and *S. cerevisiae* and applied it to CBP of lignocellulose for D-glucaric acid production. The results (Fig. 5, Table 1 and 2) are promising because it had relatively high yields, demonstrating its efficiency. However, there is still room to increase the efficiency of CBP by *T. reesei* and *S. cerevisiae*. The ratio of *T. reesei* to *S. cerevisiae* was investigated and 1:1 was the most suitable one (Fig. S5). This makes sense because decreasing the ratio weakened the lignocellulose degrading capability and increasing the ratio attenuated the fermentation by the engineered *S. cerevisiae* for D-glucaric acid production. Then we delayed the inoculation of *S. cerevisiae*, resulting reduced D-glucaric acid production. Simultaneous inoculations of *T. reesei* and *S. cerevisiae* were the best. This is because delaying *S. cerevisiae* inoculation let *T. reesei* consumed more substrates and less was left for D-glucaric acid production.

The cellulase of *T. reesei* Rut-C30 is deficient in β-glucosidase and the previous researches proved that the mixed culture of *T. reesei* and *A. niger* could improve the cellulase production and the cellulose deconstruction capability ^29, 31, 34^. Here we introduced *A. niger* into the microbial consortium and the cellulase production during CBP was enhanced (Fig. 6). On contrary, D-glucaric acid concentration lowered. It is plausible that *A. niger* got more but contributed less in the team of the three members, therefore impairing the efficiency of CBP. The same situation happened when we introduced *S. cerevisiae opi1*Δ into the microbial consortium to increase *myo*-inositol availability (Fig. 6 and Fig. S8). Then we engineered *T. reesei* Rut-C30 to make it able to provide *myo*-inositol during CBP. But once *T. reesei* was engineered, its cellulase production capability was crippled (Fig. S7). This made the microbial consortium weak in lignocellulose degradation, negatively affecting the efficiency of CBP for D-glucaric acid production from lignocellulose (Fig. 6). The failure of metabolic engineering of *T. reesei* here discourages the further efforts to metabolically engineer *T. reesei* for direct production of D-glucaric acid from lignocellulose. Even if successful, it is nearly impossible to obtain a super strain simultaneously good at cellulase production and D-glucaric acid production. Microbial cell factory would be faced with serious metabolic dilemma when doing the two things at the same time ^35^. Thus, the artificial microbial consortium of *T. reesei* Rut-C30 (cellulase specialist) and *S. cerevisiae* LGA-1 (D-glucaric acid specialist) was an excellent team with “chemistry” in CBP.

In the long run, direction production of D-glucaric acid from lignocellulose is advantageous over that from glucose, not to mention that from *myo*-inositol. Due to the highest integration of single unit operations, the utmost simplicity of bioprocess control, the cheapest substrate, and CBP is promising in the production of D-glucaric acid from lignocellulose. The relatively high D-glucaric acid titer and yield from this work proves that CBP of lignocellulose by the artificial microbial consortium of *T. reesei* and *S. cerevisiae* deserves extensive and in-depth research.

## Conclusions

The biosynthetic pathway of D-glucaric acid production was constructed in *S. cerevisiae* INVSc1 whose *opi1* was knocked out by expressing *miox4* from *A. thaliana* and *udh* from *P. syringae*, successfully obtaining two high titer D-glucaric acid producing strains, LGA-1 and LGA-C in terms of integration into Ty loci or episomal expression, respectively. Both LGA-1 and LGA-C produced record breaking titers of D-glucaric acid, indicating that *S. cerevisiae* INVSc1 was an excellent host. However, these high D-glucaric titers were facilitated by *myo*-inositol supplementation, which is not so preferable as direct production from lignocellulose. D-glucaric acid production from lignocellulose by SHF, SSF and CBP were studied here and it was found that CBP by an artificial microbial consortium of *T. reesei* Rut-C30 and the engineered *S. cerevisiae* was a promising approach with relatively high titer and yield. The microbial consortium was attempted to be designed for higher efficiency but the two members, *T. reesei* Rut-C30 and *S. cerevisiae* LGA-1, were proved to be the best teammates for CBP. Further work should be done to improve the efficiency of this microbial consortium for D-glucaric acid production from lignocellulose.

## Acknowledgements

This work was supported by the General Grant for Young Scholar (2018JQ2022) and the second-class General Postdoctoral Grant (2017BSHEDZZ100) from Shaanxi Province, the Special Funding and first-class General Financial Grants from the China Postdoctoral Science Foundation (2018T111102 and 2016M600815) and the Start-up Fund for Talent Introduction (Z111021602) from Northwest A&F University.

