## Supplementary material for "Consolidated bioprocessing of lignocellulose for production of glucaric acid by an artificial microbial consortium": Figure S1

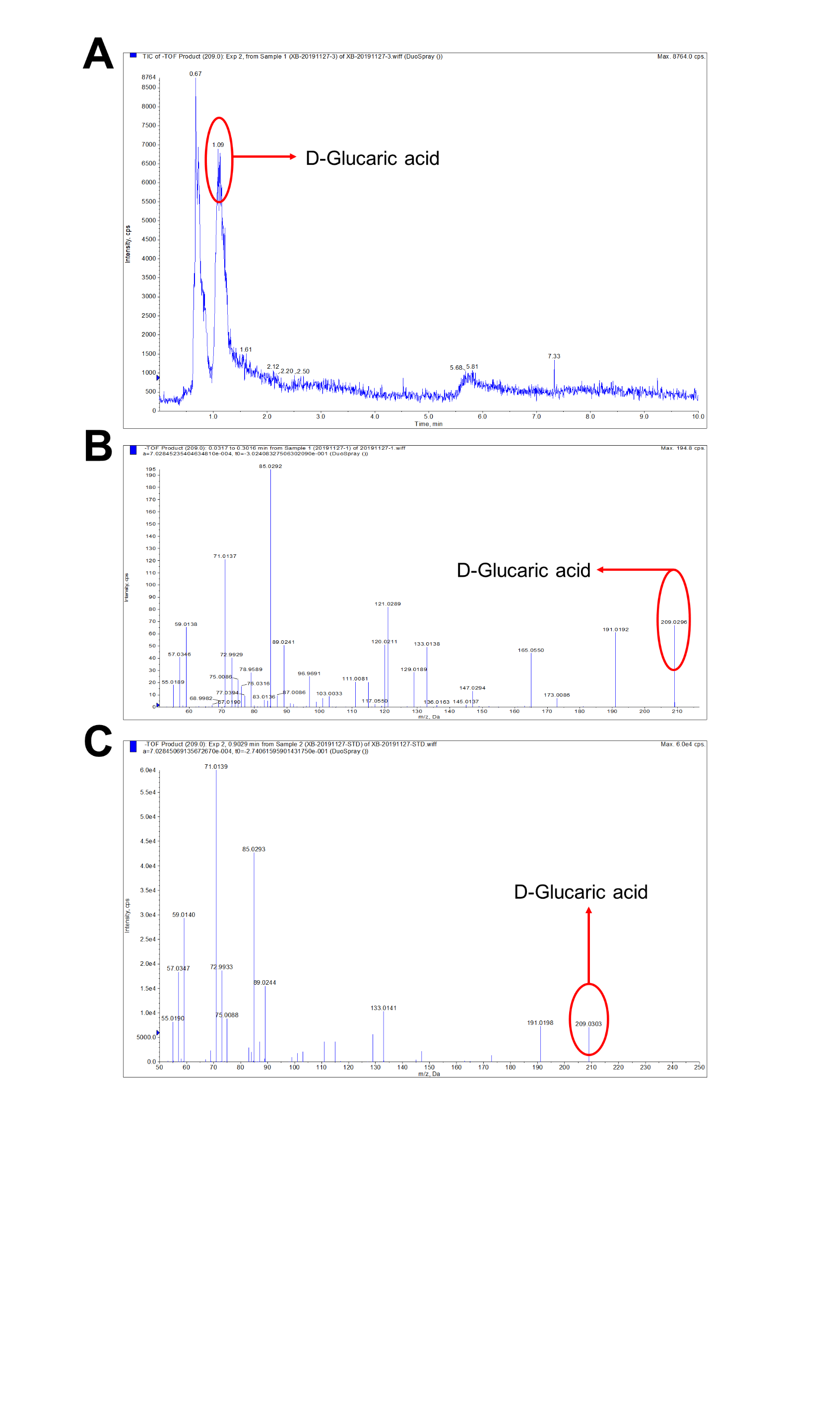

Fig. S1. Results of LC-MS analysis. LC graph (A) and MS graph (B) of the fermentation broth of *S. cerevisiae* LGA-1 after 7 d fed-batch fermentation on YPD medium supplemented with 10 g/L glucose and 10.8 g/L *myo*-inositol. (C) MS graph of the standard of D-glucaric acid.
