## Supplementary material for "Consolidated bioprocessing of lignocellulose for production of glucaric acid by an artificial microbial consortium": Figure S3

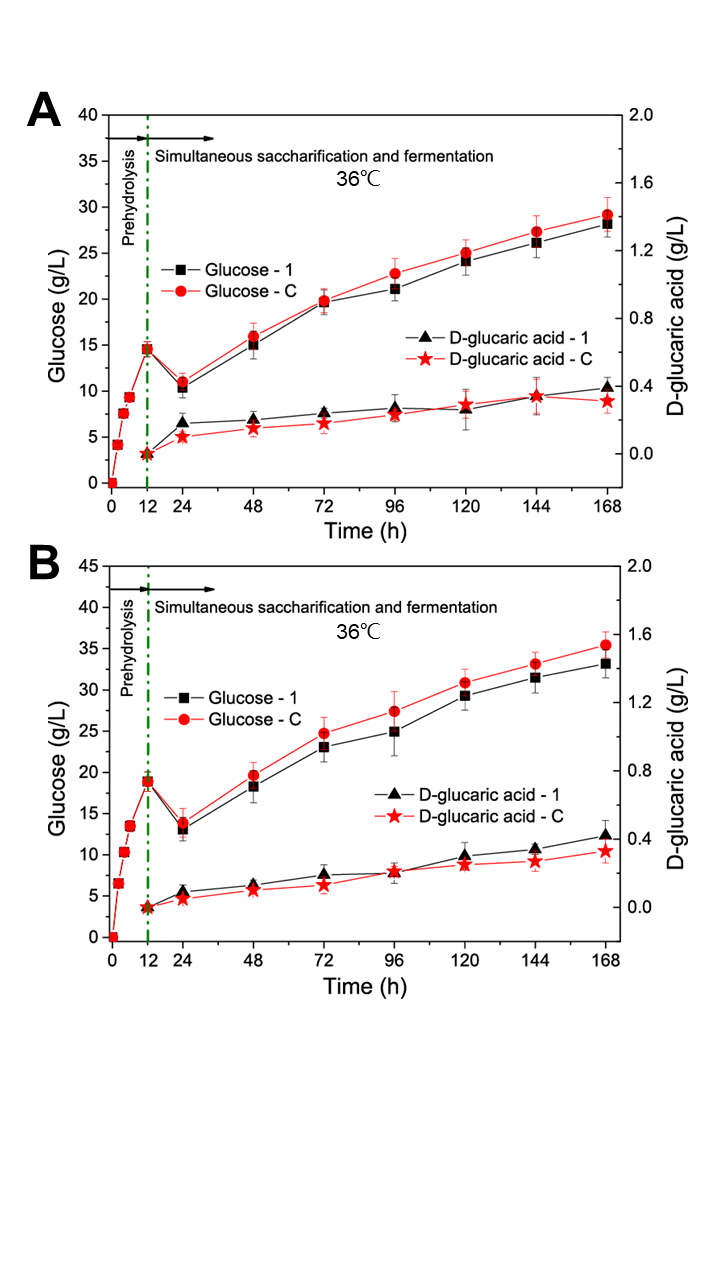

Fig. S3. Simultaneous saccharification and fermentation of Avicel (A) and steam-exploded corn stover (B) at 36℃ from 12 h to 168 h after enzymatic prehydrolysis at 50℃ for 12 h. LGA-1 and LGA-C are the engineered *S. cerevisiae* strains capable of producing D-glucaric acid. Data shown here are average values of at least three biological replicates and error bars are standard deviations.
