## Supplementary material for "Consolidated bioprocessing of lignocellulose for production of glucaric acid by an artificial microbial consortium": Figure S4

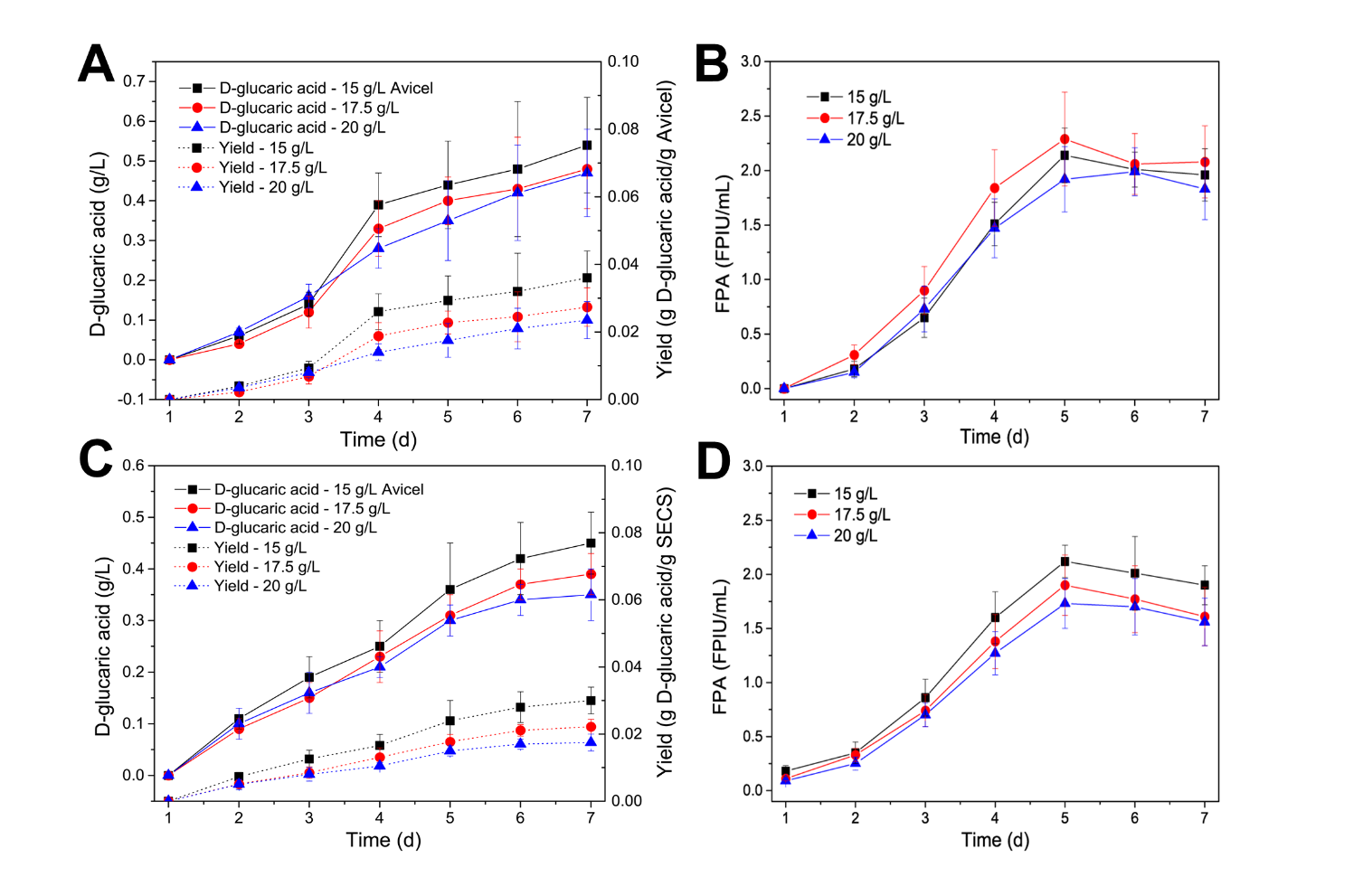

Fig. S4. Time courses of CBPs of 15, 17.5 and 20 g/L Avicel or steam-exploded corn stover (SECS) for D-glucaric acid production by *S. cerevisiae* LGA-1. (A) Concentrations of D-glucaric acid and yields during CBP of Avicel. (B) FPAs during CBP of Avicel. (C) Concentrations of D-glucaric acid and yields during CBP of SECS. (D) FPAs during CBP of SECS. Data shown here are average values of at least three biological replicates and error bars are standard deviations.
