## Supplementary material for "Consolidated bioprocessing of lignocellulose for production of glucaric acid by an artificial microbial consortium": Figure S6

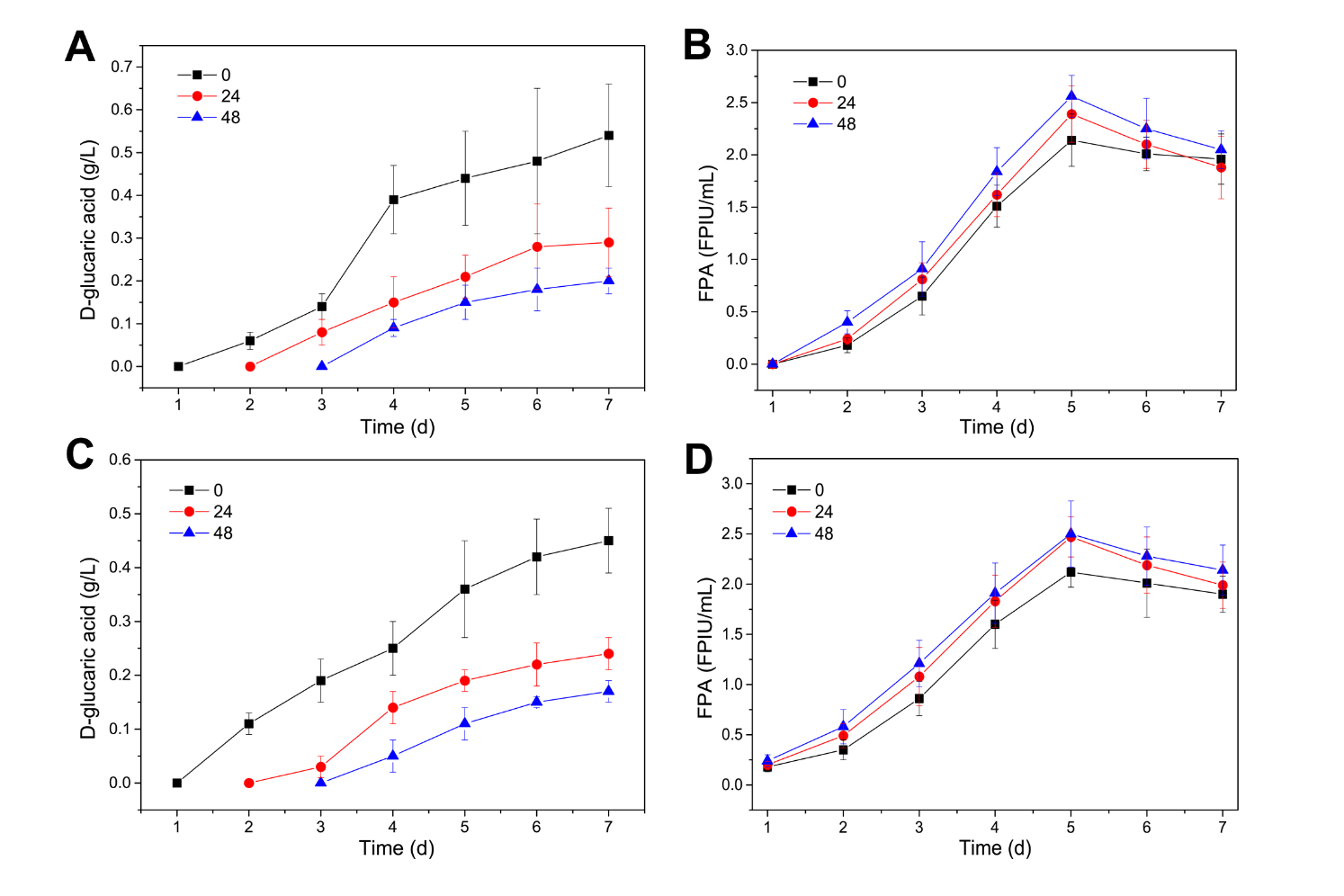

Fig. S6. Effects of the delay time of *S. cerevisiae* LGA-1 inoculation on the CBPs of 15 g/L Avicel (A and B) or steam-exploded corn stover (SECS) (C and D) by the microbial consortium of *T. reesei* Rut-C30 and *S. cerevisiae* LGA-1 for D-glucaric acid production. (A) Concentrations of D-glucaric acid during CBP of Avicel. (B) FPAs during CBP of Avicel. (C) Concentrations of D-glucaric acid during CBP of SECS. (D) FPAs during CBP of SECS. Data shown here are average values of at least three biological replicates and error bars are standard deviations.
