## Supplementary material for "Consolidated bioprocessing of lignocellulose for production of glucaric acid by an artificial microbial consortium": Figure S7

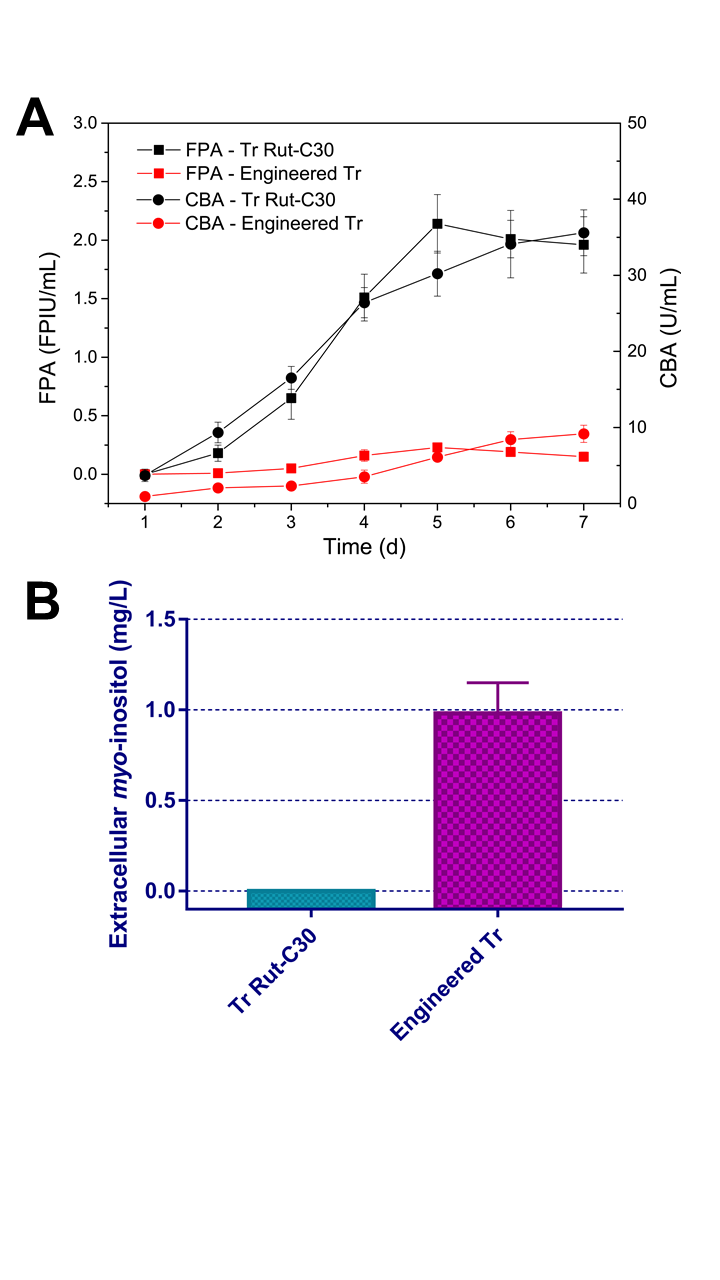

Fig. S7. (A) FPAs and CBAs during fermentation on 15 g/L Avicel by *T. reesei* Rut-C30 and the engineered *T. reesei* for *myo*-inositol production. (B) Concentrations of *myo*-inositol after 5 d fermentation. Data shown here are average values of at least three biological replicates and error bars are standard deviations.
