## Supplementary material for "Consolidated bioprocessing of lignocellulose for production of glucaric acid by an artificial microbial consortium": Figure S8

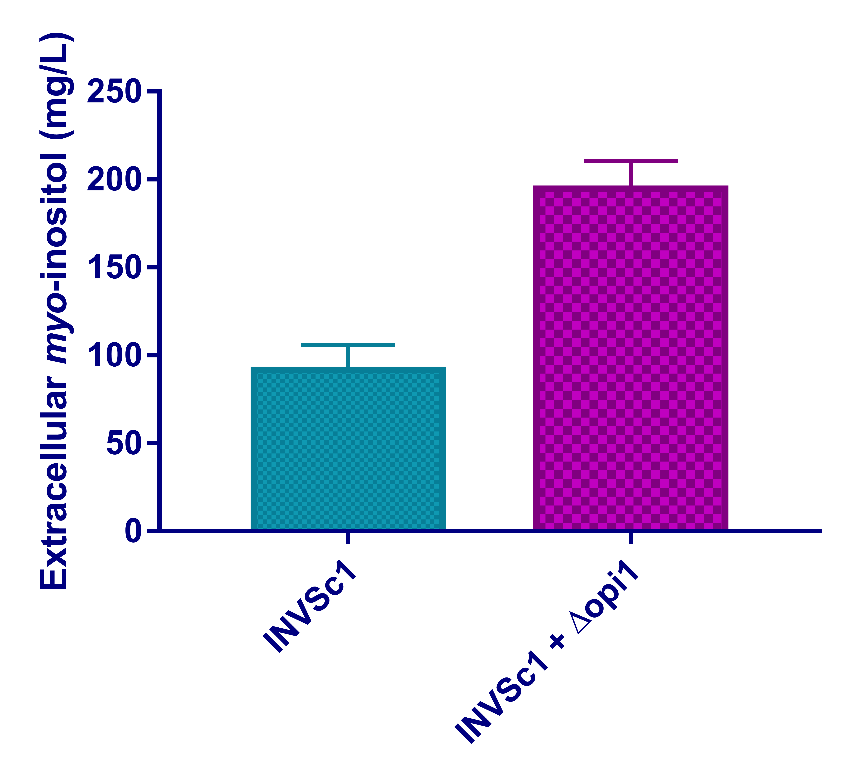

Fig. S8. Concentrations of *myo*-inositol produced by *S. cerevisiae* strains after 5 d fermentation on YPD medium. INVSc1 was the starting strain, and INVSc1 + Δopi1 the engineered *S. cerevisiae* whose *opi1* was knocked out. Data shown here are average values of at least three biological replicates and error bars are standard deviations.
