## Supplementary material for "Consolidated bioprocessing of lignocellulose for production of glucaric acid by an artificial microbial consortium": Table S1

Table S1. Plasmids and strains used in this study.

| Plasmids or strains | Descriptions | Reference |
| --- | --- | --- |
| pY26-GPD -TEF  pY26-miox4-udh  pUG6  pSH47  pCAMBIA1300  pCA-Pcbh1-ips-Tcbh1  *E.coli* DH5α  *A. tumefaciens* AGL-1  *S. cerevisiae* INVSc1  *S. cerevisiae* INVSc1 *opi1∆*  *S. cerevisiae* LGA-1  *S. cerevisiae* LGA-C  *T. reesei* Rut-C30  *A. niger* CICC2103 | under GAL regulative regulation, URA3 marker, MCS derived from pBLUESCRIPT (ColE1 (derivative) ori, f1 ori, Amp^R^)  Recombinant vector carrying *miox4* and *udh*  E. coli plasmid with segment LoxP-Kan-LoxP  Shuttle plasmid for *E. coli* and *S. cerevisiae*, Cre gene  Integrating plasmid for *T. reesei*, Shuttle plasmid for *E. coli* and *A. tumefaciences*, Kan^R^, Hyg B^R^  Recombinant vector carrying *ips*  Used for constructing plasmids  Mediating the transformation of T. reesei  MATa, His-, Leu-, Trp-, Ura-  MATa His-, Leu-, Trp-, Ura- opi1*∆0*  INVSc1 *opi1∆* carrying *miox4* and *udh*  INVSc1 *opi1∆ c*arrying pY26-miox4-udh  Used for cellulase production and starting  Used in CBP | Miaoling Bio  This study  Miaoling Bio  Miaoling Bio  Stored in author´s laboratory  This study  Stored in author´s laboratory  Stored in author´s laboratory  Stored in author´s laboratory  This study  This study  This study  Stored in author´s laboratory  Stored in author´s laboratory |
