## Supplementary material for "Consolidated bioprocessing of lignocellulose for production of glucaric acid by an artificial microbial consortium": Table S2

Table S2. Oligomers used in this study.

| Oligomer | Sequence（5’-3’） |
| --- | --- |
| knock-OPI1F | ttaaagcgtgtgtatcaggacagtgtttttaacgaagatactagtcattgCAGCTGAAGCTTCGTACGC |
| knock-OPI1R | tataatattattactggtggtaatgcatgaaagacctcaatctgtctcggGCATAGGCCACTAGTGGATCTG |
| delta1-F | GCTGGCAACTAATAGGGACAC |
| delta1-R | gaggtcgctccaattcagctGGCTATAATATCAGGTATACAGAA |
| MIOX4-F | ttctgtatacctgatattatagccAGCTGAATTGGAGCGACCTC |
| MIOX4-R | gaaataccgcacagatgcgtGCGCGCAATTAACCCTCAC |
| FURA3-F | gtgagggttaattgcgcgcACGCATCTGTGCGGTATTTC |
| FURA3-R | AAGGGCTCCCTATCTACTGGA |
| LURA3-F | CCACCCATGTCTCTTTGAGC |
| LURA3-R | tgataaactgagctcggcgcACTGAGAGTGCACCATACCAC |
| UDH-F | gtggtatggtgcactctcagtGCGCCGAGCTCAGTTTATCA |
| UDH-R | agttgatttttattccaacaTGTAAAACGACGGCCAGTGA |
| delta 2-F | tcactggccgtcgttttacaTGTTGGAATAAAAATCAACT |
| delta 2-R | AAAAGGGAATCTGCAATTCT |

The letters in lowercase are homologous sequence.
